# Constant and variable warming differentially shape bacterial coexistence through phage-mediated interactions

**DOI:** 10.64898/2026.06.14.732128

**Authors:** Zachary M. Bailey, Roxane Gualino, Claudia Bank, Madhav P. Thakur

## Abstract

Frequency-dependent predation helps maintain bacterial diversity, but its stabilizing role may be compromised under climate warming. Whereas theory suggests that warming can weaken predator-prey interactions and destabilize prey coexistence, it remains unclear whether constant and variable warming regimes differentially disrupt predation and alter coexistence outcomes in bacterial communities. Here, we experimentally tested how constant and variable warming (both +4°C above ambient, but negligible versus high thermal variance) affect the coexistence of two *Pseudomonas* species in the presence of their lytic phage. Phage predation increased competitive symmetry between the two bacterial species and promoted bacterial coexistence. Under constant warming (negligible variance), this phage-mediated frequency dependence buffered competitive asymmetries and often promoted persistence of the otherwise inferior *P. putida*, thereby magnifying coexistence. In contrast, variable warming (high variance) weakened phage control, shifted the advantage to *P. protegens*, and increased competitive exclusion events. The erosion of top-down control under variable warming was consistent with strong thermal sensitivity of phage infection rates, revealed by thermal performance curves. Our findings demonstrate that phage reduction of competitive asymmetry between bacterial hosts is amplified under constant warming but undermined under variable warming, conditions that are becoming increasingly frequent under climate change.

## Introduction

Coexistence between similar species can be maintained through a variety of mechanisms including niche partitioning, competition, and predation [1]. Theory shows that competing species can coexist when one or more of these mechanisms operate, because they prevent any single species from dominating and help maintain a stable ecological state [2, 3]. Predation is one of such commonly studied mechanisms, which has been shown to both promote and destabilize prey coexistence depending on the abiotic environments [4–6]. Predation can promote coexistence between two species when they predate more on the superior competitor either through prey preference or density dependent predation [1, 4]. By contrast, predation can erode coexistence when it preferentially targets the inferior competitor or acts independently of prey dominance, thereby weakening negative frequency dependence and accelerating competitive exclusion [4, 7]. Moreover, predation-mediated coexistence is environmentally contingent; for instance, abiotic stress such as temperature can alter predator density and foraging traits [6, 8, 9], thereby weakening top-down regulation of prey populations and destabilizing coexistence [7].

Ongoing climate change, such as warming, is altering predator-prey interactions with detrimental consequences for predator-mediated species coexistence and ecological stability [7, 10, 11]. Increasing temperatures due to climate change encompass both constant increases in average temperatures, and erratic fluctuations in temperature, such as through periodic heat waves [12–17]. These two types of temperature shifts, constant and variable, lead to different outcomes partially based on species traits. For instance, variable temperatures often mimic real-world scenarios more and affect life history traits such as development and survival rates more than constant temperature [16, 18]. Constant warming, while less representative of natural thermal regimes, allows for controlled isolation of mean temperature effects on biological processes such as growth, metabolism, and species interactions [19–23]. However, warming experiments that exclude thermal variability may misrepresent real-world biological responses and generate misleading predictions about species interactions and community dynamics [19, 24, 25]. Microbial systems are uniquely suited to test how constant versus variable warming alters predator-prey interactions, offering precise environmental control, large population sizes, and sufficient replication to resolve community-level outcomes [26–29].

Here, we utilized a two species bacterial prey system of *Pseudomonas protegens* and *Pseudomonas putida,* each with a specialized bacteriophage predator, φGP100 and Psp1, respectively. Bacteriophages or phages are the obligatory viruses of bacteria, infecting bacteria to replicate and killing them in the process [30, 31]. Although phages are obligate intracellular parasites, lytic phages such as φGP100 and Psp1, impose top-down, frequency dependent mortality on their bacterial hosts while producing new viral particles. This consumer-like interaction is analogous to predation and motivates our utilization of predator-prey terminology, which, has a long history in microbial ecology [32, 33]. Phages provide immense ecological importance as they tend to liberate large amounts of organic carbon from bacteria each day [34, 35]. This viral shunt is an important ecological process for cycling nutrients in the biosphere [36–39]. Lytic phages with narrow host ranges allow for bacterial coexistence by killing members of the most dominant bacterial species the “kill the winner” hypothesis [40, 41]. The ability of phages to enable coexistence is dependent on the co-evolution between bacteriophages and their bacterial prey where bacteria that gain phages resistance can overcome phage pressure [42–44]. *Pseudomonas protegens* is a strong competitor against many other soil bacteria especially other *Pseudomonas spp.*[45]; they can produce a variety of secondary metabolites with antimicrobial capabilities limiting the growth and survival of neighbouring bacterial species [45–47]. In contrast, *Pseudomonas putida* lacks the offensive capabilities of *P. protegens*, but can grow on a wide variety of substrates including aromatic compounds [48, 49].

Using *P. protegens* and *P. putida* as two prey species, along with their lytic bacteriophages φGP100 and Psp1, we assessed how constant and variable warming alter phage-mediated bacterial coexistence. Specifically, we tested two sets of hypotheses (Figure 1): (1) Because φGP100 preferentially infects the dominant competitor *P. protegens*, its presence will reduce the competitive asymmetry between species and promote coexistence, whereas Psp1, which infects the inferior competitor *P. putida*, will accelerate competitive exclusion. (2) Variable warming will weaken φGP100 -mediated coexistence, whereas constant warming will strengthen the same phage mediated coexistence (Figure 1).

**Figure 1:**
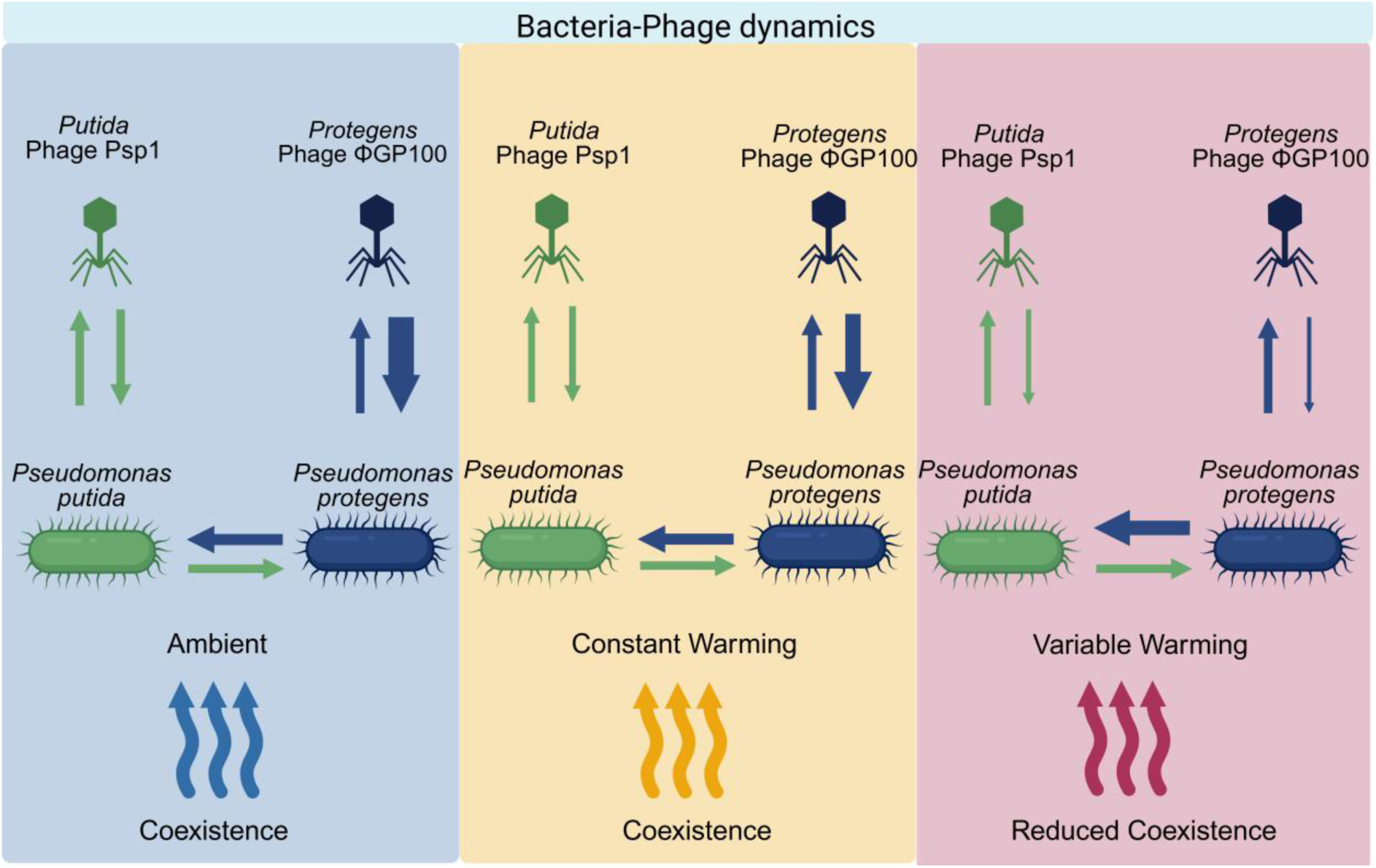
A schematic showing potential interactions between members of our microbial community in three different warming treatments (ambient, constant and variable warming). Predicted coexistence outcome between two bacterial species, *Pseudomonas protegens* and *Pseudomonas putida* are shown through horizontal straight arrows. Phage-bacteria interactions are shown by vertical straight lines. The thickness of arrows indicates the strength of interactions between species (at three different warming regimes).

## Methods

### Species and culture conditions

We used two bacterial species in our experiments, *Pseudomonas protegens* CHAO and *Pseudomonas putida* MM1. We used two phage species, φGP100 and Psp1, that are specialized predators infecting *P. protegens* or *P. putida* respectively, but are unable to actively infect the other species [48, 50, 51]. All bacteria were grown in M9 minimal media made using 100 ml of 5x M9 minimal salts with 4 ml of 20% Glucose (W/V), 6 ml of 20% Arabinose (W/V), 1 ml of MgSO_4_7H_2_O (1M), 50 µl of CaCl_2_ (1M) and then filled to 500 ml with milliQ water. This formulation, hereafter M9 60:40, was used in all experiments because differential arabinose utilization by *P. putida* (but not *P. protegens*) supports their stable coexistence (Supplementary Fig 1). We used Sodium Magnesium (SMG) buffer for our dilutions with 5.8 g NaCl, 2 g MgSO_4_·7H_2_O, 50 mL 1M Tris HCl, (pH 7.5), 5 ml of 2% gelatine solution (filter sterilized) and milliQ water to 1 L total volume. Populations of both *P. protegens* and *P. putida* were grown up as overnight cultures at 28°C, shaking at 220 RPM in 4 ml of M9 media 60:40. These overnight cultures were then re-inoculated at 28°C in 20 mL of M9 60:40.

### Experimental design

We established a two-species bacterial community composed of the inferior competitor *P. putida* and the superior competitor *P. protegens*. These bacteria were exposed to specialized phage predators, *Psp1* (for *P. putida*) and φGP100 (for *P. protegens*). We conducted a 9-day serial transfer experiment under three temperature regimes: (1) ambient, reflecting local summer conditions (details below); (2) constant warming (ambient + 4°C); and (3) variable warming (ambient + 4°C with stochastic fluctuations generated using a sinusoidal model to simulate diurnal temperature variability). The ambient temperature profiles were derived from MeteoSwiss records for Bern (46.993° N, 7.467° E) [52], Switzerland (2013–2023 averages from June, July and August). The constant and variable warming treatments shared the same mean temperature but differed in temporal structure: constant warming applied a uniform +4°C offset, while variable warming introduced fluctuations around the daily mean, varying the magnitude of warming across each day (Supplementary Fig. 2). The +4°C offset reflects end-of-century warming projected for Switzerland under high-emission scenarios [53, 54]. These temperature regimes were programmed into Percival incubators (Model E-36L2, Percival Scientific Inc., Perry, IA, USA) using the built-in ramping function to simulate diurnal thermal fluctuations. These temperatures were also measured in real time using a HOBO Pendant MX2201 Bluetooth data logger to record the temperature every five minutes. Each combination of bacteria and phages was tested in a full factorial design with six replicates across all biotic and temperature treatments, including blanks that only contain media to monitor for bacterial cross-contamination.

Populations were cultivated in 96 deep-well plates (2 ml) and transferred after 24 hours of growth into a new 96 deep-well plate with 1ml of fresh media per each well (1:100). We used 100 µl of culture mixed with 100 µl of 50% glycerol to make freezer stocks at -80° C each day. Bacterial abundance was quantified by flow cytometry (Invitrogen Attune CytPix with CytKick Autosampler; Thermo Fisher Scientific, USA) on Days 1, 3, 5, 7, and 9 post-inoculations. The flow cytometry outputs were first size gated with the SSC-A (Side Scatter Area) and the SSC-H (Side Scatter Height) to filter debris and doublets followed by gating using the FSC-A (Forward Scatter Area) and fluorescent detector channel BL1-A (488 nm). We measured the cell counts of both bacterial species using the flowCore package version 2.20 [55] in R statistical software [56].

### Thermal performance curves

To understand the mechanisms underlying phage-mediated bacterial interactions under different warming regimes, we characterized thermal performance curves for all species used in the serial transfer experiment. The temperature gradient used to generate performance curves spanned the range experienced across the three experimental treatments (12–40°C).

For the measurements of phage thermal performance curves, we used the ancestral bacterial species for the bacterial lawns and grew them on LB agar plates with LB overlay agar. LB agar being made with 20 g of LB Medium (Lennox) from Carl Roth (Art. No. X964.2) and 15g of Agar-Agar, Kobe I Carl Roth (Art. No. 5210.2) for 1 L of LB agar. LB overlay agar is made with 20 g of LB Medium (Lennox) from Carl Roth (Art. No. X964.2) and 7g of Agar-Agar, Kobe I Carl Roth (Art. No. 5210.2) for 1 L of LB overlay agar.

Phage stocks of Psp1 and φGP100 were standardized to ∼10^8^ PFUs/mL and the overnight *P. protegens* and *P. putida* were re-inoculated and grown at 220 RPM and 28°C until reaching an OD_600_ of ∼1 (∼10^7^ CFUs/mL) and are in exponential growth phase. Eppendorf tubes are used with 1 ml of bacterial culture (∼10^7^ CFUs/ml) and 10 µl of phage stock (∼10^6^ PFUs/mL) with three replicates of each bacteria-phage combination at each temperature. We measured seven different temperatures for each set of bacteria and phages to include the entire range of thermal conditions of our experiment where the lowest recorded temperature was 15.96° C and the highest was 35.95°C. These temperatures were 12°, 16°, 20°, 24°, 28°, 32°, 36°, and 40° C. Phage-bacteria culture combinations were incubated in Percival E-36L2 incubators for four hours at their set temperature. After incubation, 100 µl of chloroform was added to each tube, three tubes of the initial phage-bacteria inoculum were also chloroformed. These tubes were vortexed and centrifuged at 10,000 x g for 5 minutes separating the aqueous phase (containing phages) from the chloroform and cell debris.

20 µl from the aqueous phase was taken from each tube to make a dilution series. Phages were plated on overlay agar plates and incubated at 28° C overnight. We then counted the plaques of the initial phages and phages grown over four hours of inoculation calculating PFUs as Plaques × Dilution Factor. The growth rates of our bacteriophages were calculated using equation (1) where *N_4_* is the PFUs after 4 hours and *N_0_* is the initial PFUs.

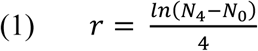

For bacterial thermal performance curves, each species was grown in overnight cultures at 28°C with shaking at 220 RPM that was then re-inoculated into 96 microwell plates with a dilution of 1:100. We used bacterial cultures from ancestral bacteria as well as from day 5 and day nine clones to evaluate the bacterial population’s acclimatization to heat via their thermal performance curves. We also examined each bacterial population with and without their specialized phage to measure the effects of phages on population growth. We read the OD600 values of our microplates over 24 hours at 20°, 24°, 28°, 32°, 36° and 40° C using a Biotek Synergy H1 microplate reader (Agilent, Santa Clara, California, United States) every 10 minutes. We then analyse these growth curves in R to get the growth rate (r) and area under the curve (AUC) as measures of bacterial growth that were then used in our analysis [56].

## Data analysis

### Serial transfer experiment

All analysis was done in R (v 4.5.0) [56]. Mixed effect models were implemented in R using *glmmTMB* package (v 1.1.13) [57, 58], Effsize [59] DHARMa [60], and emmeans [61]. Our fixed effects were warming regimes with three levels (ambient, + 4°C (constant), + 4°C (variable)), predation with four levels (no predator present, specialized predator, predator of other bacteria, both predators), competition with two levels (competitor, no competitor), and experimental days with two levels (three versus nine). We chose these two days as comparisons as day three marked a sufficient acclimatization time and further showed a local maximum for population sizes for some treatments (time series data shown in Supplementary figure 3), whereas day nine represented the end of the experiment (Supplementary figure 3). The well position, i.e., where each population was located within the plate, was treated as a random factor within our model. Cohen’s *d* effect sizes were calculated for the population response of both bacterial species across all treatment combinations using the Effsize package [59]. Linear model assumptions (normality of residuals and homogeneity of variance) were assessed visually using diagnostic plots with DHARMa [60], and response variables were log-transformed when these assumptions were violated.

### Thermal performance curves

Thermal performance curves were analyzed using the *growthrates* package in R (v 0.8.5) [62] for growth measures such as growth rates, and the Area Under the Curve (AUC). The thermal performance curves for bacteria and phages were calculated with the *rTPC* package (v 1.0.4) [63] and the *nls.multstart* package (v 2.0.0) [64] using growth rates calculated from the *growthrates* package for bacteria and the manually calculated growth rate for phages following the methods explained in Padfield et. al. (2021).

## Results

### Interactive effects of warming regimes, predation and competition on bacteria and phages

Both constant and variable warming altered bacterial densities relative to ambient temperatures (Table 1, Fig. 1). The presence and absence of phage predation also affected bacterial abundance (Table 1, Fig. 1). Moreover, we found a significant three-way interaction among predation, competition and warming regimes affecting bacterial abundance (Fig. 1, Table 1). Across treatments, variable warming caused stronger declines in bacterial densities than constant warming, particularly for *P. putida*. Under variable warming and in both the absence and presence of phages, *P. putida* densities were significantly reduced relative to ambient and constant-warming (Fig. 2a, Supplementary Fig. 3a). By contrast, *P. protegens* population densities varied little across warming regimes and tended to survive at any temperature (Fig. 2b, Supplementary Fig. 3b).

**Figure 2:**
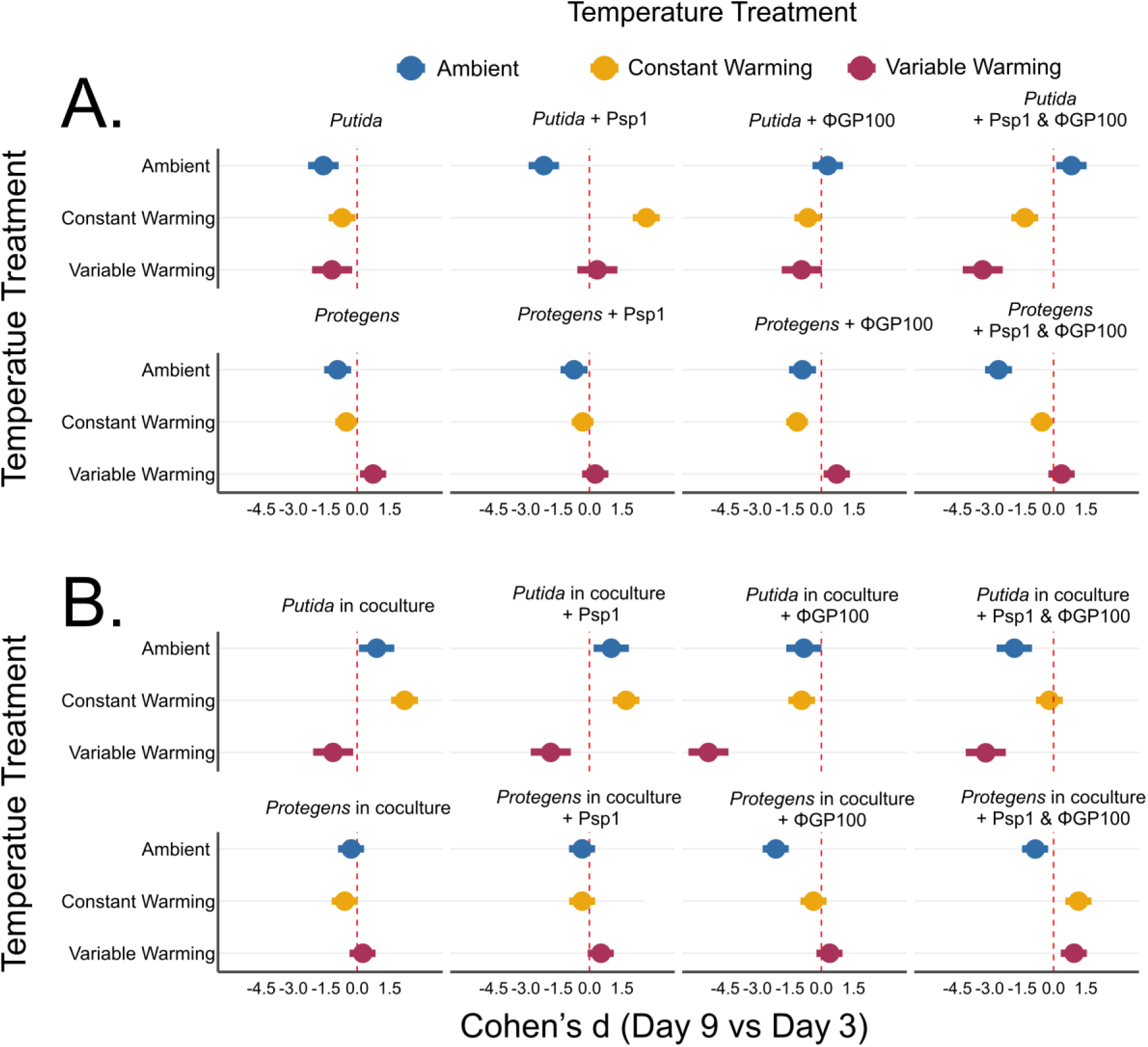
Bacterial density shifts under three different warming regimes over the course of experiment. (A) The Cohen’s *d* effect sizes (mean ±95% confidence intervals) obtained from the mixed effect models comparing bacterial densities of *Pseudomonas putida* (*putida* for brevity) and *Pseudomonas protegens* (*protegens* for brevity) **monocultures** between Day 3 and Day 9 across treatments. (B) The Cohen’s *d* effect sizes (mean ±95% confidence intervals) obtained from the mixed effect models comparing bacterial densities of *Pseudomonas putida* (*putida* for brevity) and *Pseudomonas protegens* (*protegens* for brevity) **co-cultures** between Day 3 and Day 9 across treatments.

**Table 1.** Fixed-effect tests from generalized linear mixed-effects models of bacterial abundance. Type III Wald χ² tests are shown for generalized linear mixed-effects models evaluating the effects of warming regime, competition, predation, day, and their interactions on log-transformed bacterial abundance. Separate models were fitted for *Pseudomonas putida* and *Pseudomonas protegens*. Fixed effects included warming regime (W), competition (C), predation (P), day (D), and all interaction terms, with well included as a random intercept. The table reports the fixed-effect terms, Wald χ² statistic with degrees of freedom as subscripts, associated *P* value, and Species.

| Predictors | Chi-Squared | p-value | Species |
| --- | --- | --- | --- |
| Intercept | 212.064<sub>1</sub> | **4.87E-48** | putida |
| Warming regimes (W) | 0.945<sub>2</sub> | 0.623 | putida |
| Competition (C) | 85.194<sub>1</sub> | **2.70E-20** | putida |
| Predation (P) | 23.499<sub>3</sub> | **3.18E-05** | putida |
| Day (D) | 6.905<sub>1</sub> | **0.0086** | putida |
| W×C | 29.859<sub>2</sub> | **3.28E-07** | putida |
| W×P | 26.481<sub>6</sub> | **0.000181** | putida |
| C×P | 50.108<sub>3</sub> | **7.58E-11** | putida |
| W×D | 1.793<sub>2</sub> | 0.408 | putida |
| C×D | 8.556<sub>1</sub> | **0.00344** | putida |
| P×D | 16.083<sub>3</sub> | **0.00109** | putida |
| W×C×P | 30.955<sub>6</sub> | **2.59E-05** | putida |
| W×C×D | 5.862<sub>2</sub> | 0.0533 | putida |
| W×P×D | 56.999<sub>6</sub> | **1.83E-10** | putida |
| C×P×D | 31.493<sub>3</sub> | **6.69E-07** | putida |
| W×C×P×D | 36.083<sub>6</sub> | **2.66E-06** | putida |
| Intercept | 383.683<sub>1</sub> | **1.96E-85** | protegens |
| Warming regimes (W) | 0.396<sub>2</sub> | 0.82 | protegens |
| Competition (C) | 0.072<sub>1</sub> | 0.789 | protegens |
| Predation (P) | 10.811<sub>3</sub> | 0.0128 | protegens |
| Day (D) | 5.035<sub>1</sub> | 0.0248 | protegens |
| W×C | 0.519<sub>2</sub> | 0.771 | protegens |
| W×P | 10.511<sub>6</sub> | 0.105 | protegens |
| C×P | 3.28<sub>3</sub> | 0.35 | protegens |
| W×D | 6.853<sub>2</sub> | 0.0325 | protegens |
| C×D | 1.219<sub>1</sub> | 0.27 | protegens |
| P×D | 16.173<sub>3</sub> | **0.00105** | protegens |
| W×C×P | 5.058<sub>6</sub> | 0.536 | protegens |
| W×C×D | 1.361<sub>2</sub> | 0.506 | protegens |
| T×P×D | 13.486<sub>6</sub> | 0.0359 | protegens |
| C×P×D | 14.57<sub>3</sub> | **0.00222** | protegens |
| W×C×P×D | 8.91<sub>6</sub> | 0.179 | protegens |

Phage presence in general increased the coexistence between two bacterial species, although only in ambient and constant warming regimes (Fig. 2a, 2b). However, this phage-mediated coexistence disrupted under variable warming (Fig. 2, Supplementary Fig. 3). Mixed-effects models revealed that *P. putida* experienced the largest density reductions when exposed simultaneously to variable warming, competition with *P. protegens*, and predation by their specialized predator ΦGP100 (Fig. 2a). These combined stressors frequently resulted in population extinction of *P. putida* (Supplementary Fig. 3a, Supplementary Fig 4a), whereas all *P. protegens* populations persisted throughout the experiment (Supplementary Fig. 3b, Supplementary Fig. 4b). The response of *P. protegens* to its phage ΦGP100 depended strongly on temperature regime. Under ambient conditions, ΦGP100 significantly reduced *P. protegens* densities. Under variable warming, however, *P. protegens* densities increased in the presence of ΦGP100 relative to ambient (Fig. 2b), indicating reduced phage efficacy at higher temperatures in the variable warming regime.

### Thermal performance curves

We measured the thermal performance of our phage predators and their bacterial prey with temperatures that cover the entire range of our experiment (Fig. 3). *Protegens* phage GP100 and putida phage Psp1 both increase in their growth rate until reaching a T_opt_ of ∼26°C with a T_max_ at ∼28°C (Fig 3a). When measuring the temperature within the incubator we find that the ambient and constant warming regimes did not exceed the max temperature of either phage (Fig 3b). Utilizing growth rate estimates, we find that the thermal optimum of our bacteria is generally much higher than that of their phages. The ancestral *P. putida* had a T_opt_ of 34.4 °C without phages and 32.4 °C with phages. Ancestral *P. protegens* had a T_opt_ of 37.8 °C without phages and 36.9 with phages. By the end of the experiment, *P. putida* clones from day nine had a T_opt_ of 37.9 °C in variable warming conditions without phages and 30.4 °C with phages (Supplementary Fig. 5). Similarly, *P. protegens* clones from day nine had a T_opt_ of 38.2 °C without phages and 27.1 °C with phages (Supplementary Fig. 5). Similarly, the cumulative growth estimates from our area under the curve (AUC) data suggest that ancestral *P. putida* has a T_opt_ of 33.75 °C without phages and ancestral *P. protegens* has a T_opt_ of 34.88 °C (Supplementary Fig. 6). While on day nine, *P. putida* had a T_opt_ of 31.4 without phages and 33.5 with phages, and *P. protegens* was 31.69 °C without phages and 25.54 with phages (Supplementary Fig. 6).

**Figure 3:**
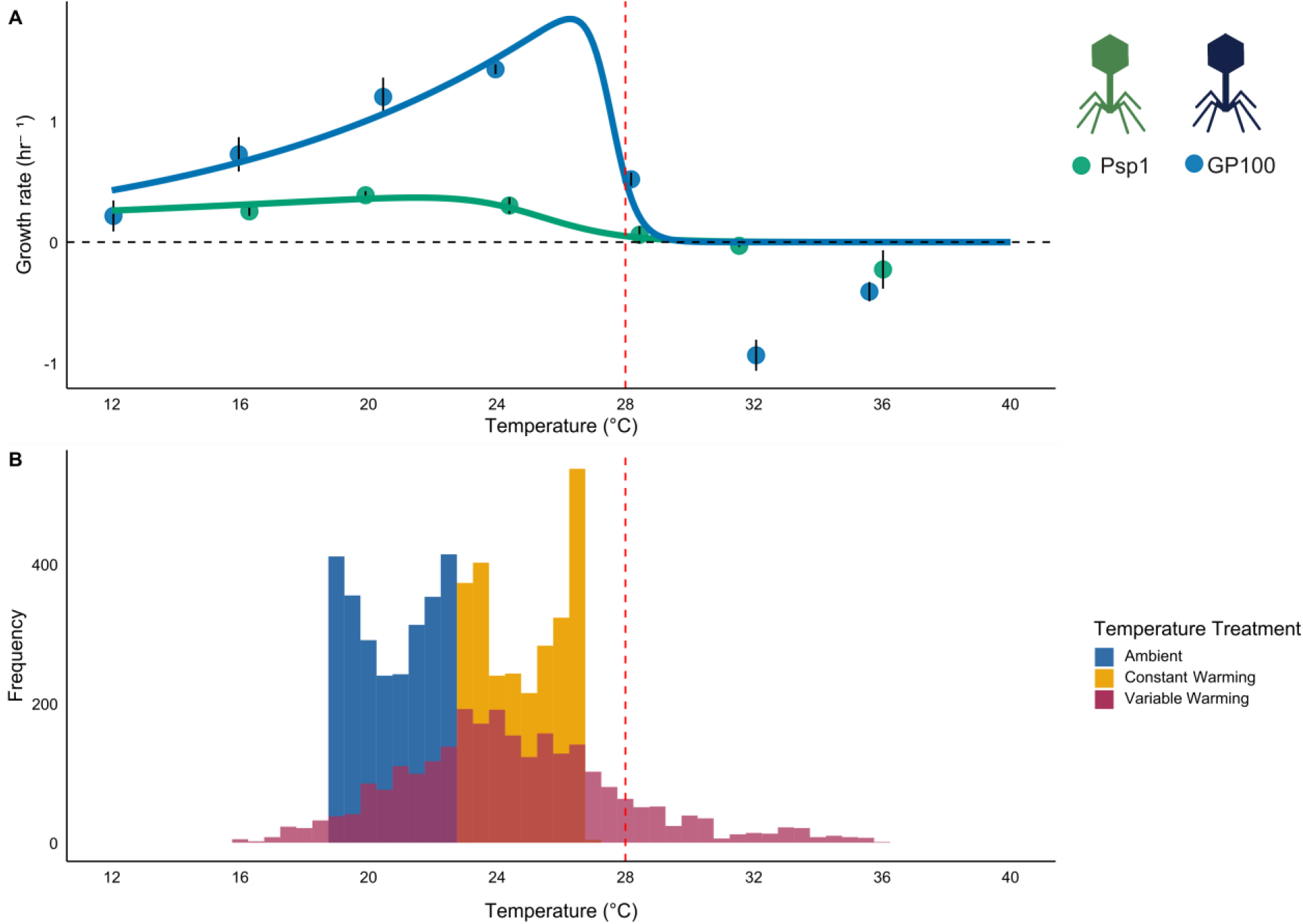
Thermal performance curves and temperature frequency in three warming regimes. (A) Thermal performance curve of both the *P. putida* phage Psp1 and the *P. protegens* phage phiGP100 demonstrating the average growth at eight different temperatures spanning the thermal regimes utilized in the serial transfer experiment. (B) Temperature distributions of the recorded temperatures within Percival incubators across the three warming regimes over all nine days of the experiment. Temperature was recorded in five-minute increments, and frequency is the number of these increments at a specific temperature.

## Discussion

Climate change is altering not only mean environmental temperatures but also the frequency, magnitude, and the temporal structure of thermal extremes [12–14, 16, 66]. Whereas constant warming experiments have provided substantial insight into how temperature influences microbial growth and interactions, far less is known about how thermal variability affects community stability and coexistence [67–70], particularly in systems regulated by antagonistic interactions such as phage predation [71, 72]. To our knowledge, this is the first experimental demonstration that constant and variable warming regimes produce markedly different outcomes for phage-mediated bacterial coexistence, despite sharing the same mean temperature. These differences emerge through temperature-dependent changes in phage performance that alter competitive interactions between bacterial hosts, compounded by shifts in bacterial performance itself. Our findings demonstrate that highly variable warming can destabilize top-down control, thereby amplifying competitive asymmetries between two competing bacterial species.

### Variable warming weakens phage-mediated stabilization of bacterial coexistence

Under ambient conditions, reciprocal phage predation promoted coexistence between *P. protegens* and *P. putida*, consistent with classic expectations of top-down regulation of prey diversity [30, 73]. However, this stabilizing effect was contingent on temperature regime. Under variable warming, phage regulation weakened substantially, leading to reduced bacterial coexistence and increased extinction risk for *P. putida*. In contrast, constant warming preserved phage-mediated coexistence despite equivalent mean temperatures. These results indicate that short-term thermal excursions, rather than elevated temperature per se, can disrupt predator–prey coupling in microbial systems [24, 74–77]. Such effects are unlikely to be captured by constant warming experiments and may therefore represent an underappreciated pathway by which climate change alters microbial interaction networks [25, 28, 78].

### Asymmetric thermal responses drive competitive exclusion

The loss of coexistence under variable warming was primarily driven by asymmetric thermal responses among bacteria and their phages. Phages alter the thermal performance of their hosts, and bacteriophages often exhibit mismatches between their thermal performance and the thermal performance of their host [79]. *Pseudomonas putida* experienced pronounced density reductions under variable temperature regimes, particularly in the presence of *P. protegens* (Fig. 2, Supp Fig. 3, Supp Fig. 4.). In contrast, *P. protegens* persisted across all treatments, highlighting differences in thermal tolerance and competitive ability. Importantly, these outcomes were mediated by temperature-dependent changes in phage efficacy. Phage replication declined sharply at elevated and variable temperatures, consistent with the observed reduction in top-down control. Phages typically respond to extreme temperature variance with either a lower absorption rate or changes to the bacterial cell structure reducing phage infections [80, 81]. Once a phage is inside a bacterial cell, thermal stress reduces the efficacy of the bacterial cellular machinery that phages utilize for replication [82]. Additionally, bacteria under stress tend to have a slower growth rate giving phage defence systems more time to become active [83]. As phage pressure relaxed, *P. protegens* was released from predation and able to competitively exclude *P. putida*. These findings support the hypothesis that thermal mismatches between predators and prey can shift interaction outcomes and destabilize coexistence [84, 85], even over relatively short ecological timescales [72, 79]. This is relevant as phage driven effects are generally temporally limited as phage resistance easily evolves when phages and bacteria are in co-culture for extended periods of time [42, 51, 86].

## Conclusions

Microbial communities underpin ecosystem processes ranging from biogeochemical cycling to host health, and their stability is often maintained by antagonistic interactions including competition and phage predation [43, 68, 71]. Our results suggest that increasing thermal variability may systematically weaken these interactions, favouring thermally resilient and competitively dominant taxa. This shift could reduce microbial diversity and alter community function under future climate scenarios. This contrasts with constant warming where increases in temperature that remain constant still allow for coexistence. Temperature increases that do not extend past the thermal tolerability and remain predictable can still be acclimatized to. More broadly, our study highlights the need to incorporate thermal variance and extremes into experimental and theoretical frameworks used to predict microbial responses to climate change. By demonstrating that warming regime structure can override mean temperature effects, our findings caution against extrapolating from constant warming experiments alone. Accounting for thermal variability will be essential for accurately forecasting how climate change reshapes microbial coexistence and the ecological processes it governs.

## Author contributions

**ZMB**: Conceptualization, Methodology, Investigation, Formal Analysis, Writing – Original Draft.

**RG**: Investigation, Writing – Review & Editing.

**CB:** Methodology (supporting), Writing – Review & Editing

**MPT**: Conceptualization, Methodology (supporting), Resources, Funding Acquisition, Writing – Original Draft (supporting), Writing – Review & Editing.

## Acknowledgement

We thank Anine Wyser for her help in the experiment. This project was funded by the Swiss National Science Foundation (Grant 310030_212550 for MPT).

