## Supplemental Figures for "Constant and variable warming differentially shape bacterial coexistence through phage-mediated interactions"

### Supplementary Figures

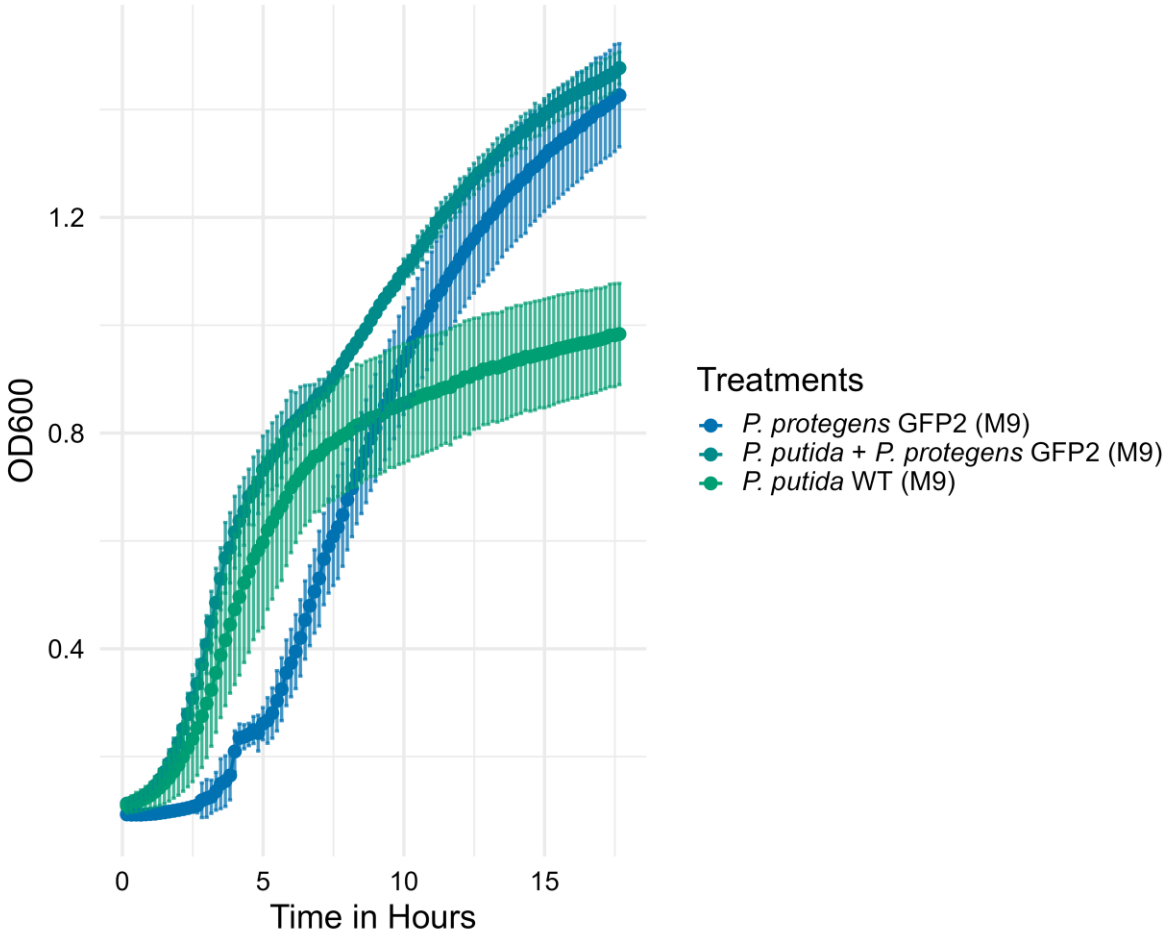


**Supplementary Figure 1**: Growth curves over 16h of both monocultures and co-cultures of *Pseudomonas protegens* and *Pseudomonas putida* using M9 60:40 media, a combination of arabinose and glucose with M9.  Co-cultures tend to have a bi-phasic growth following the rapid increase of *P. putida* followed by *P. protegens*.

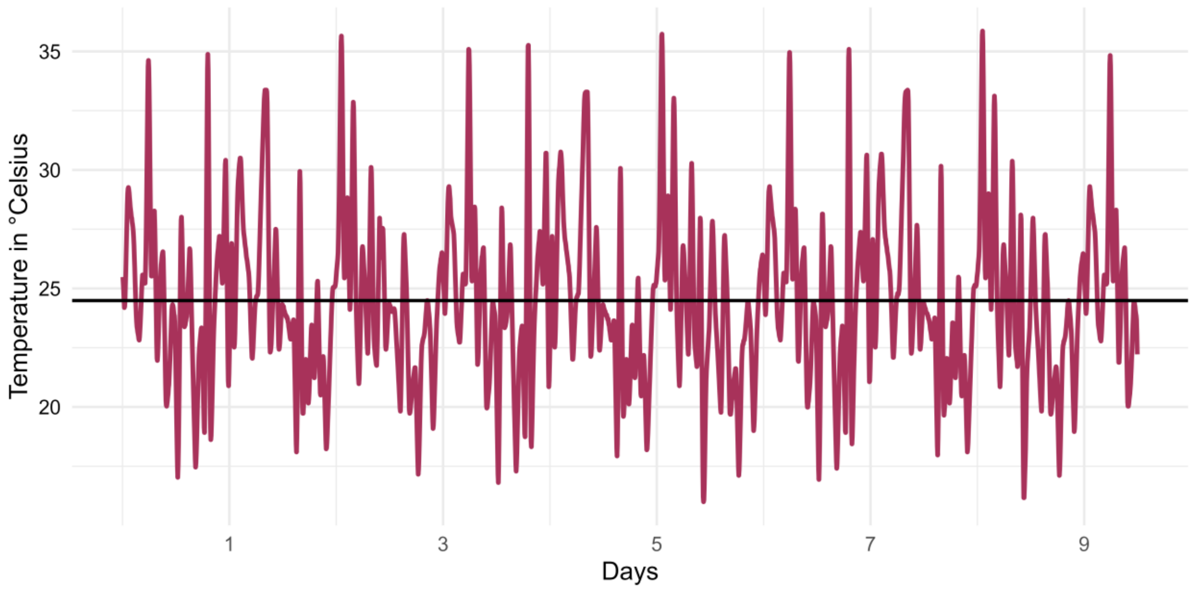


**Supplementary Figure 2:** The red line denotes the recorded temperature within the variable warming regime Percival incubator over the course of the experiment. The mean daily temperature is indicated by a solid black line. The variable temperature regime has three distinct sub-regimes that were each repeated three times.

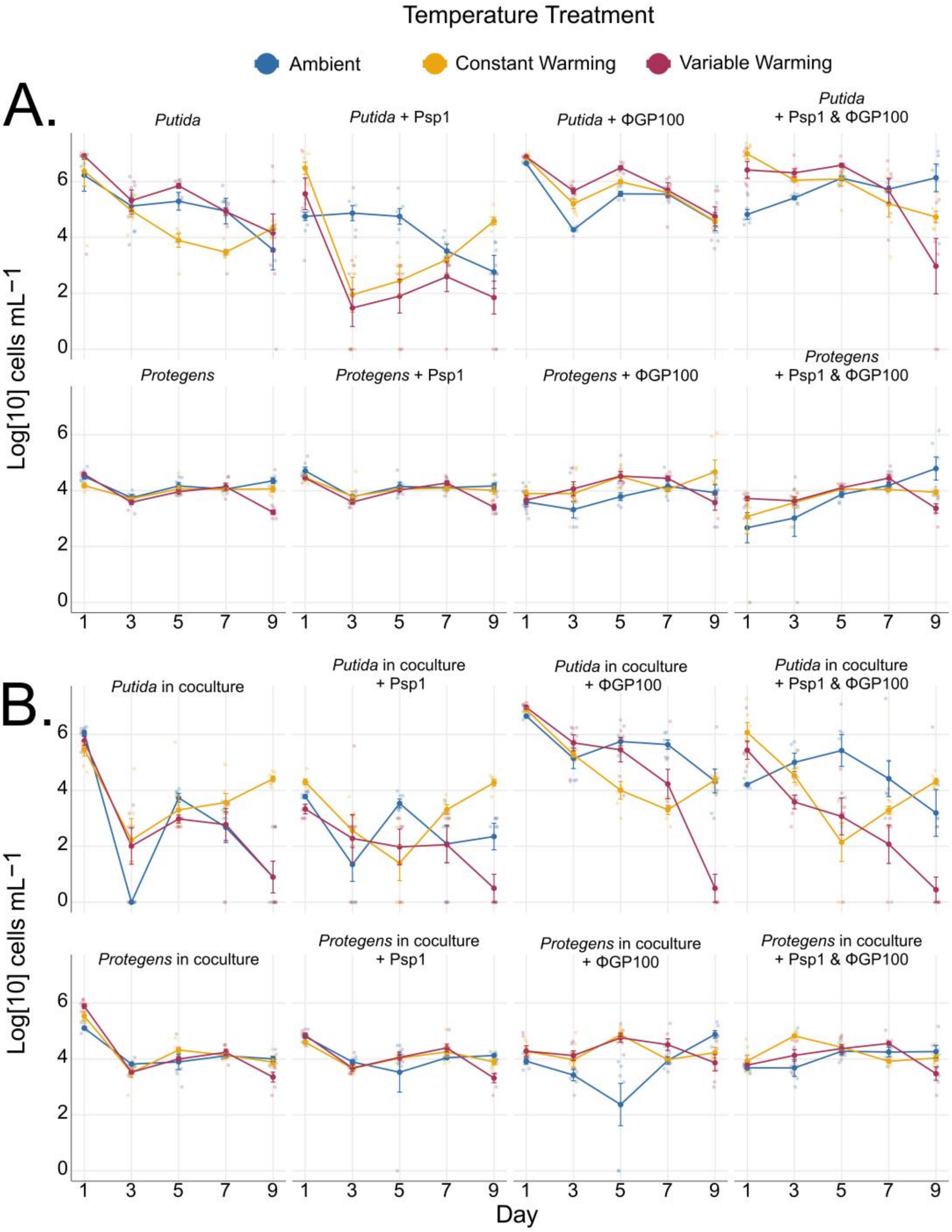


**Supplementary Figure 3:** (A) Bacterial cell counts log-transformed from the flow cytometry of *Pseudomonas putida* and *Pseudomonas protegens* in monocultures across the 9 days of our experiment. The first row is all four *P. putida* combinations, and the second row has all four monoculture combinations of *P. protegens*. (B) Bacterial cell counts log-transformed from the flow cytometry of *P. putida* and *P. protegens* in coculture across the nine days of our experiment. The first row is the measurement of *P. putida* cell counts where all four combinations are *P. putida* cells in coculture with *P. protegens*, and the second row is *P. protegens* cell counts where all four combinations are *P. protegens* in coculture with *P. putida*.

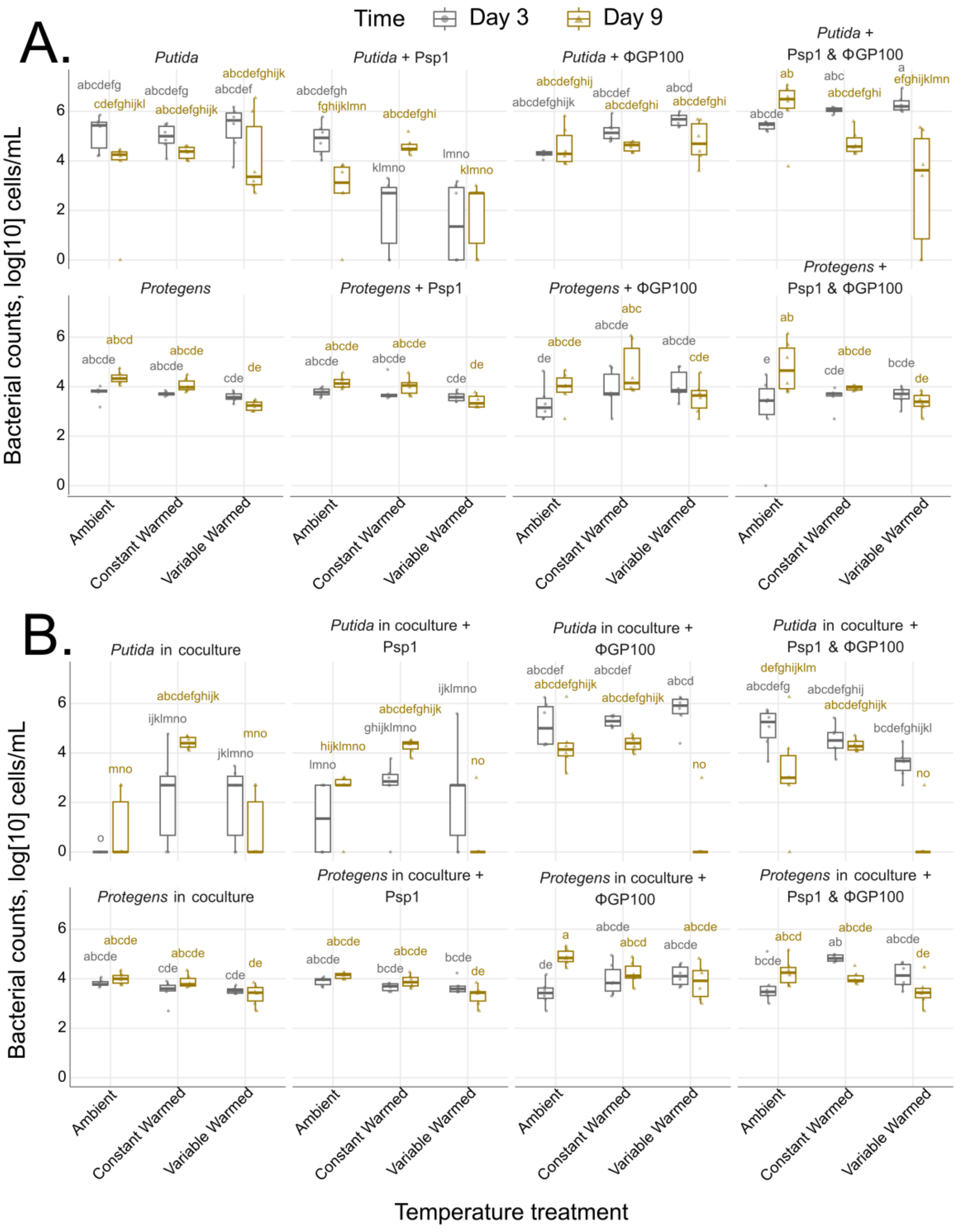


**Supplementary Figure 4:** (A) *Pseudomonas putida* cell counts from days 3 and 9 of our serial transfer experiment where letters indicate significant differences between groups based on ANOVA followed by Tukey's HSD test, using results from the three-way interaction (warming treatment, microbial communities and time) in our linear models. (B) *Pseudomonas protegens* cell counts from days three and nine of our serial transfer experiment where letters indicate significant differences between groups based on ANOVA followed by Tukey's HSD, using results from the three-way interaction (warming treatment, microbial communities and time) in our linear models.

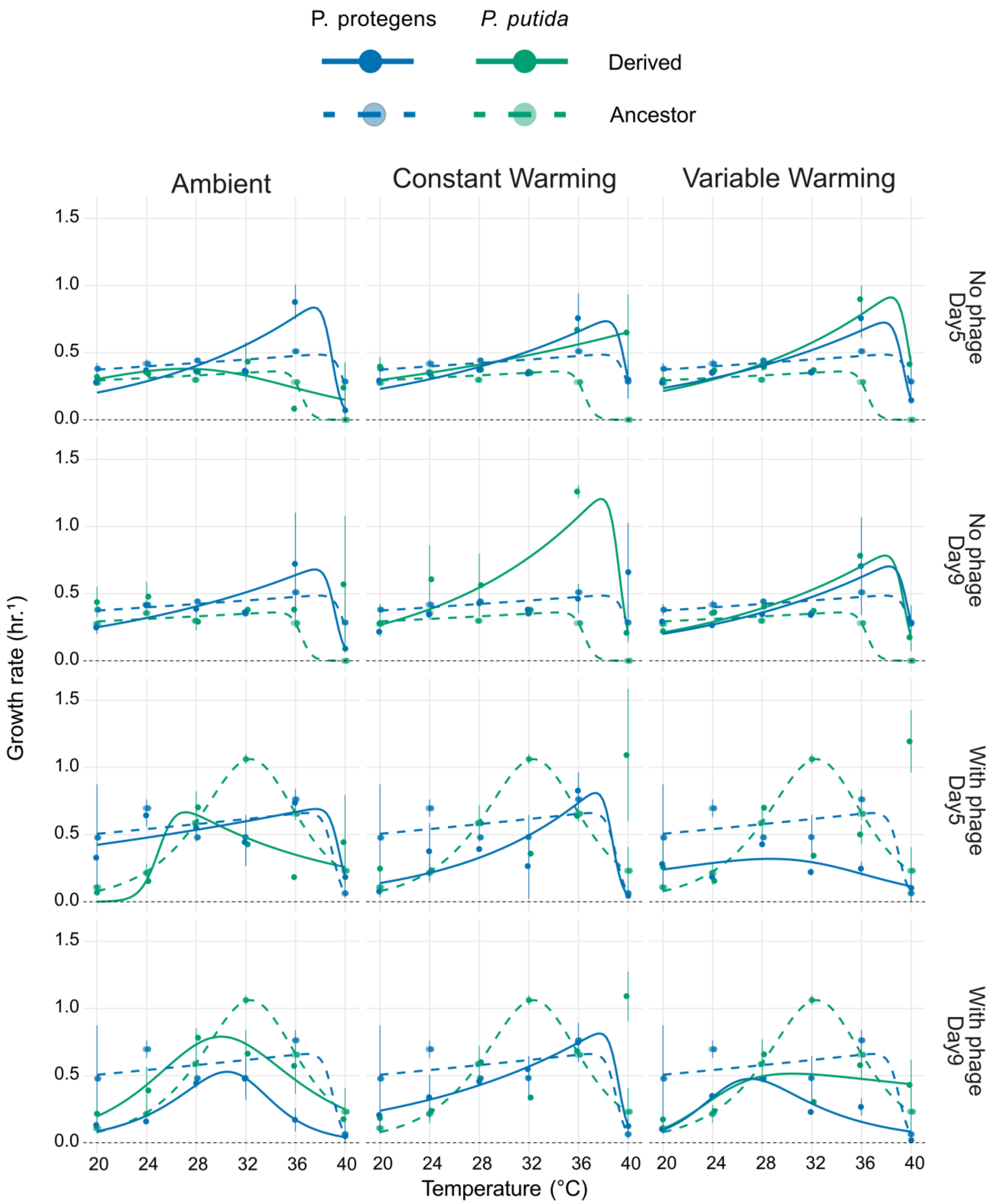


**Supplementary Figure 5:** TPC of bacteria populations utilizing the growth rate estimates of *growthcurver* from Day 5 and Day 9 (solid lines) in contrast to their ancestral, Day 0, (dashed line) counterparts. This compares the growth at different temperatures of *P. putida* (green) and *P. protegens* (blue) both with and without their respective phage predator, Psp1 for *P. putida* and ΦGP100 for *P. protegens*.

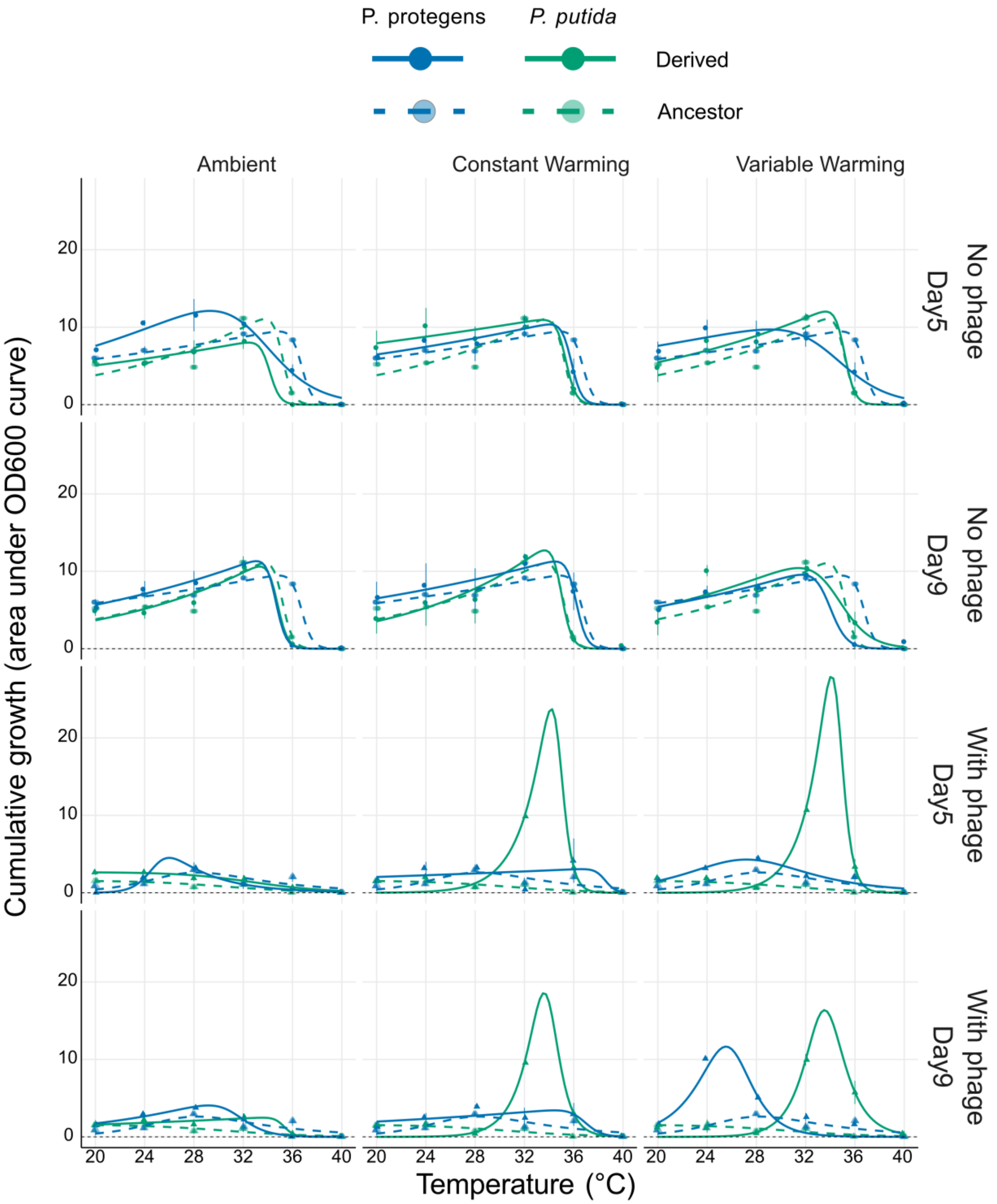


**Supplementary Figure 6:** TPC of bacteria populations utilizing the area under the curve (AUC) cumulative growth estimates of *growthcurver* from Day 5 and Day 9 (solid lines) in contrast to their ancestral, Day 0, (dashed line) counterparts. This compares the growth at different temperatures of *P. putida* (green) and *P. protegens* (blue) both with and without their respective phage predator, Psp1 for *P. putida* and ΦGP100 for *P. protegens*.
